# Optimization of Seizure Prevention by Cannabidiol (CBD)

**DOI:** 10.1101/2024.08.24.609518

**Authors:** Bidhan Bhandari, Sahar Emami Naeini, Sholeh Rezaee, Hannah M Rogers, Hesam Khodadadi, Asamoah Bosomtwi, Mohammad Seyyedi, Neil J MacKinnon, Krishnan M Dhandapani, Évila Lopes Salles, David C Hess, Jack C Yu, Debra Moore-Hill, Fernando L. Vale, Lei P Wang, Babak Baban

## Abstract

**Objective:** Cannabidiol (CBD) is one of the most prominent non-psychotropic cannabinoids with known therapeutic potentials. Based on its anti-seizure efficacy, the first cannabis derived, pharmaceutical grade CBD-based medication was approved in the USA in 2018 for the treatment of seizures in patients 2 years and older. Despite the effectiveness in reducing seizures, there remain several major questions on the optimization of CBD therapy for epilepsy such as the optimal dosage, composition, and route of delivery, which are the main objective of this current study.

**Methods:** We evaluated the antiseizure effects of CBD through different compositions, routes of delivery, and dosages in a pre-clinical model. We used a kainic acid-induced epilepsy model in C57BL/6 mice, treated them with placebo and/or CBD through inhalation, oral and injection routes. We used CBD broad spectrum (inhaled and injection) versus CBD isolate formulations. We employed the Racine scaling system to evaluate the severity of the seizures, flow cytometry for measuring Immune biomarkers and neurotrophic factors, and histologic analysis to examine and compare the groups.

**Results:** Our findings showed that all forms of CBD reduced seizures severity. Among the combination of CBD tested. CBD broad spectrum via inhalation was the most effective in the treatment of epileptic seizures (p<0.05) compared to other forms of CBD treatments.

**Conclusion:** Our data suggest that route and CBD formulations affect its efficacy in the prevention of epileptic seizures. Inhaled broad spectrum CBD showed a potential superior effect compared to other delivery routes and CBD formulations in the prevention of epileptic seizures, warrants further research.

## Introduction

Cannabinoids are closely related naturally occurring compounds found in *Cannabis sativa*. The cannabis plant has been used for its medicinal purposes since ancient times all over the world (1). However, it was only in the last half century and particularly for last two decades that the public and scientific interests have mounted significantly, triggering intense and extensive research in systematic medical characterization of cannabinoids (1). The cannabis plant produces over 500 compounds, but only 100 of them are classified as cannabinoids, and two of them, delta-9-tetrahydrocannabinol (THC) and Cannabidiol (CBD) being the most known and investigated cannabinoids for their medical as well as recreational purposes (2-3). While THC is recognized and well-known for its psychoactive features, CBD is non-psychotropic, with potential anti-inflammatory effects, and a high therapeutic index (4). Recent work by our laboratory and others suggests beneficial effects of CBD alone or in combination with other cannabinoids in the treatment of several pathologic conditions including neurologic diseases as well as malignancies (5-8). Importantly, several reports have suggested that CBD could be used as an anti-convulsive and well-tolerated agent with beneficial effects in the treatment of seizures (9-10). It was in 2018 when the US Food and Drug Administration (FDA) approved the first pharmaceutical dosage form of CBD for treating patients 2 years and older with Dravet syndrome (DS) or Lennox-Gastaut syndrome (LGS), two of the various, rare epileptic disorders classified as epileptic encephalopathies (10-11). The treatment later received European Commission approval to be available for patients across Europe (12).

Epilepsy is a common and chronic neurologic condition characterized by recurrent and wanton seizures (13). Affecting over 50 million individuals at all ages globally, epilepsy remains one of the most challenging disorders with direct, indirect, and intangible costs to individuals, healthcare system and the economy overall, such as worker productivity (13-15). One of the major difficulties in managing epilepsy relates to existing therapeutic modalities (16-18). Most of the current Anti-Seizure Medications (ASMs) are inconsistent in their efficacy and, furthermore, they are associated with undesirable side effects (17-19). In fact, because most of the current research on ASMs focus only on controlling the epileptic seizures rather than treating the underlying causes, a significant (30%) number of epileptic patients are classified as pharmacoresistant (non-responsive to treatment) (18-19). Pharmacoresistant epileptic patients are characterized with refractory epilepsy, requiring polytherapeutic regiments which may cause drug-drug interactions, adding to the complexity of treating drug resistant epilepsy (18-20). Furthermore, the chronic nature of pharmacoresistant epilepsy makes the evaluation of adverse effects of ASMs very challenging in long-term trials (20). Such therapy-induced involution along with the complex nature of epilepsy, warrants the exploration of new therapies to treat epilepsy more effectively and safely based on more definitive, conclusive evidence.

Since FDA approval of CBD-based therapy for epilepsy, there has been no optimization research addressing many questions concerning the best route of administration, efficacy, and ideal dosage in the treatment of refractory epileptic seizures (21-22). In addition, the cost of has raised concerns about its affordability by patients and health insurers as well as for further investigational purposes (23-24).

In this study, we have investigated the prophylactic, prevention of epileptic seizure using inhalant CBD in a murine model. To our knowledge, this is the first study of its kind in this field. Our findings describe the efficacy of our preparation not only in the prevention of epileptic seizures, but also showing a promising superior effectiveness of the dosage form when compared to other CBD-based formulations in the prevention of epileptic seizures.

## Materials and methods

### Animals

Male C57BL/6 mice purchased from Jackson Laboratories USA were used in these experiments. Mice were housed as a group and handled according to the National Institute of Health (NIH) guide for the Care and Use of Laboratory Animals. All experiments were conducted under the approval of the Augusta University Animal Care and Use Committee (Protocol # 2011-0062).

### Experimental groups and prophylactic treatments with variable delivery routes

Mice were divided into 6 groups (n = 5/group, total of 5 independent cohorts). The “CBD” group which was treated prophylactically with inhalant CBD (8.5 mg/mouse) using inhaled CBD in three different doses of CBD concentration of 100%, 10% and 1% (TM Global Bioscience USA). The formulation of experimental inhalers was a broad-spectrum CBD containing natural compounds and cannabinoids (except THC. The placebo was administered through inhalers too. However, for the placebo, the whole CBD content was replaced by hemp seed oil, contained no cannabinoids. As depicted in figure 1, the inhaler was modified by adding an extra nozzle piece to adjust to the mouse model and to further control the intake of CBD. The second, “i. p.” group, received CBD (broad-spectrum) prophylactically through intra-peritoneal route (Liquid 1000 mg) while the third group (Placebo) received inhaled placebo. The fourth group was treated with CBD isolate through oral gavage (Epidiolex, NDC70127-100-60, Cardinal Health). The CBD isolate formulation contained CBD only with no other cannabinoids and compounds, which differs it from our inhaled formulation. After 30 minutes of CBD/Placebo treatments, the epileptic seizures were induced in all mice using Kainic acid as described hereinafter.

**Figure 1.**
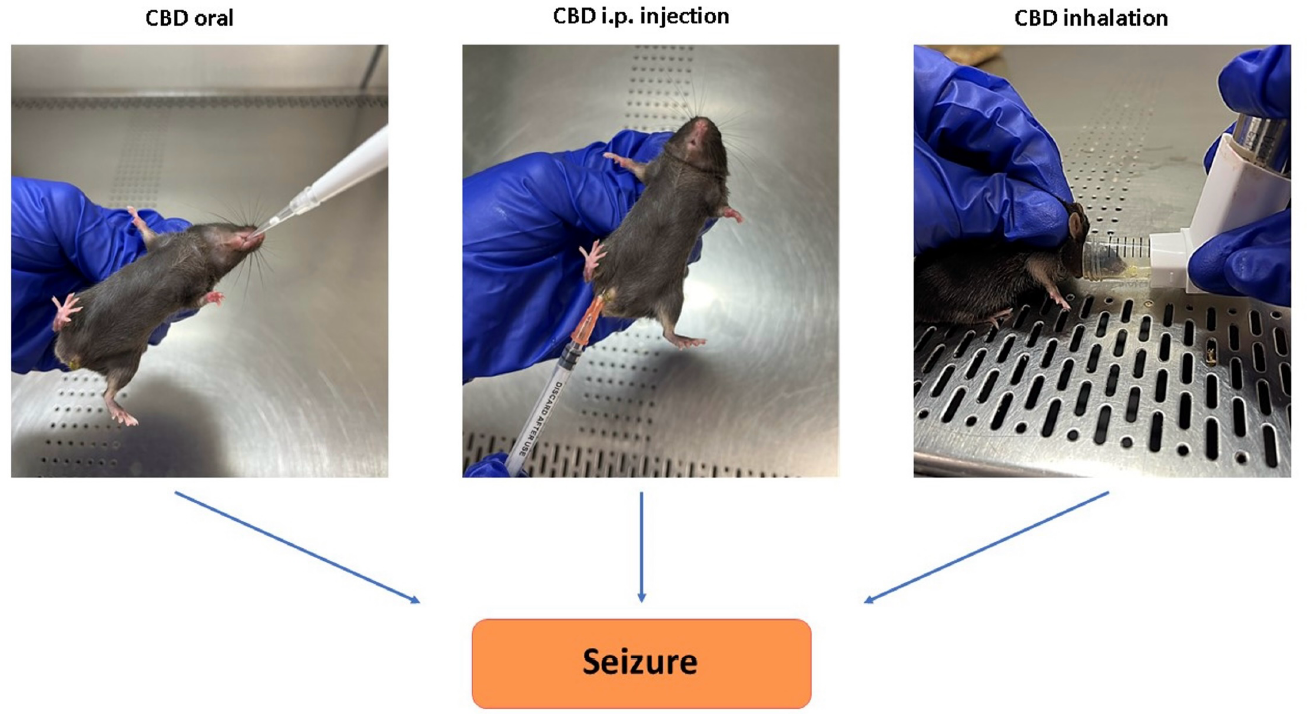
Graphical display of variable methods used for CBD delivery. a) Modified Inhaler along with actual delivery of CBD through inhalation., b) CBD administration through oral route, simulating form of delivery of CBD-based treatment as used in clinical practice for the treatment of epileptic seizures., C) CBD administration through intraperitoneal injections (i. p.).

### Induction of Epileptic Seizures: Kainic acid infusion

Kainic Acid (Fisher Scientific) was used to induce epileptic seizures as described previously (25). Briefly, Kainic Acid (KA) was dissolved in phosphate buffered saline (pH 7.4) and delivered through intra-peritoneal injection (20 mg/kg). The Racine Scoring system was used to assess the behavior and severity of the induced epileptic seizures.

### The Racine Scoring System

The Racine scoring system was used to assess the degree of seizure in our experimental epilepsy model. The scoring was based on the standard Racine scale as described previously (26-27). Briefly, the level of seizure severity was measured with the following stages: (0), no abnormality; (1), Mouth and facial movements; (2) Head nodding; (3) Forelimb clonus; (4) Rearing; (5) Rearing and falling. While the scoring was performed and monitored in real time, however, the whole process was video recorded for further and future analysis if needed.

### Analytical Flow cytometry

To set a pattern of potential predictive, diagnostic, and prognostic biomarkers during the course of study, whole blood and brain samples were collected, processed and analyzed using flow cytometry. Both brain tissues and whole blood were analyzed for immune-inflammatory biomarkers including PD1 (Programmed cell death protein 1, an inhibitory receptor), Interleukin 33, Interleukin 6 (IL-33/IL-6, alarmin/cytokine), and level of Brain-derived neurotrophic factor (BDNF).

### Preparative flow cytometry

For cell-surface marker of Programmed cell death protein 1 (PD-1), immunophenotyping was carried out as described previously (Khodadadi et al; 2021). Briefly, for brain samples, tissues were sieved through a cell strainer (BD Biosciences), followed by centrifugation (252 g, 5 min, 4°C) to prepare single-cell suspensions. Next, all samples (whole blood and cell suspension from brain) were incubated with conjugated antibody against PD-1, followed by fixation and permeabilization using fix/perm concentrate (eBioScience). Next, samples were incubated with antibodies for intracellular staining of IL-33, IL-6 and BDNF (all antibodies were purchased from Biolegend unless otherwise noted). Cells were then run through a NovoCyte Quanteun flow cytometer. Cells were gated based on forward and side scatter properties and on marker combinations to select viable cells of interest. Single stains were used to set compensation, and isotype controls were used to determine the level of nonspecific binding. Analysis was performed using FlowJo (version 11.0) analytical software. Cells expressing a specific marker were reported as a percentage of the number of gated events. A population was considered positive for a specific marker if the population exceeded a 2% isotypic control threshold.

### Statistics

For statistical analysis, data were analyzed using Graphpad Prism 9. A one-way analysis of variance (ANOVA) followed by Kruskal-Wallis multiple comparisons test was performed to establish significance (p < 0.05) among all groups in the RACINE score analysis while in the flow cytometry analysis the one-way ANOVA was followed by Tukey’s test.

## Results

### Prophylactic treatment with Inhaled CBD was able to effectively mitigate severity of KA-induced acute epileptic seizures

Treatment of mice with the inhalant CBD in a prophylactic fashion (30 min prior to seizure induction) reduced the severity of seizures significantly compared to placebo treatment. The seizure severity was measured using the Racine seizure scale from 0-5 as described in methods and shown in figure 2A. While mice treated with three different concentration (100%, 10% and 1%) of inhaled CBD were scored between1-2, the placebo treated mice were scored at 4.5 with full spectrum of severe seizure (Fig 2A-B, p<0.05). Among all three concentrations of CBD, the full strength CBD showed the most effective outcomes with significant differences (p<0.01).

**Figure 2.**
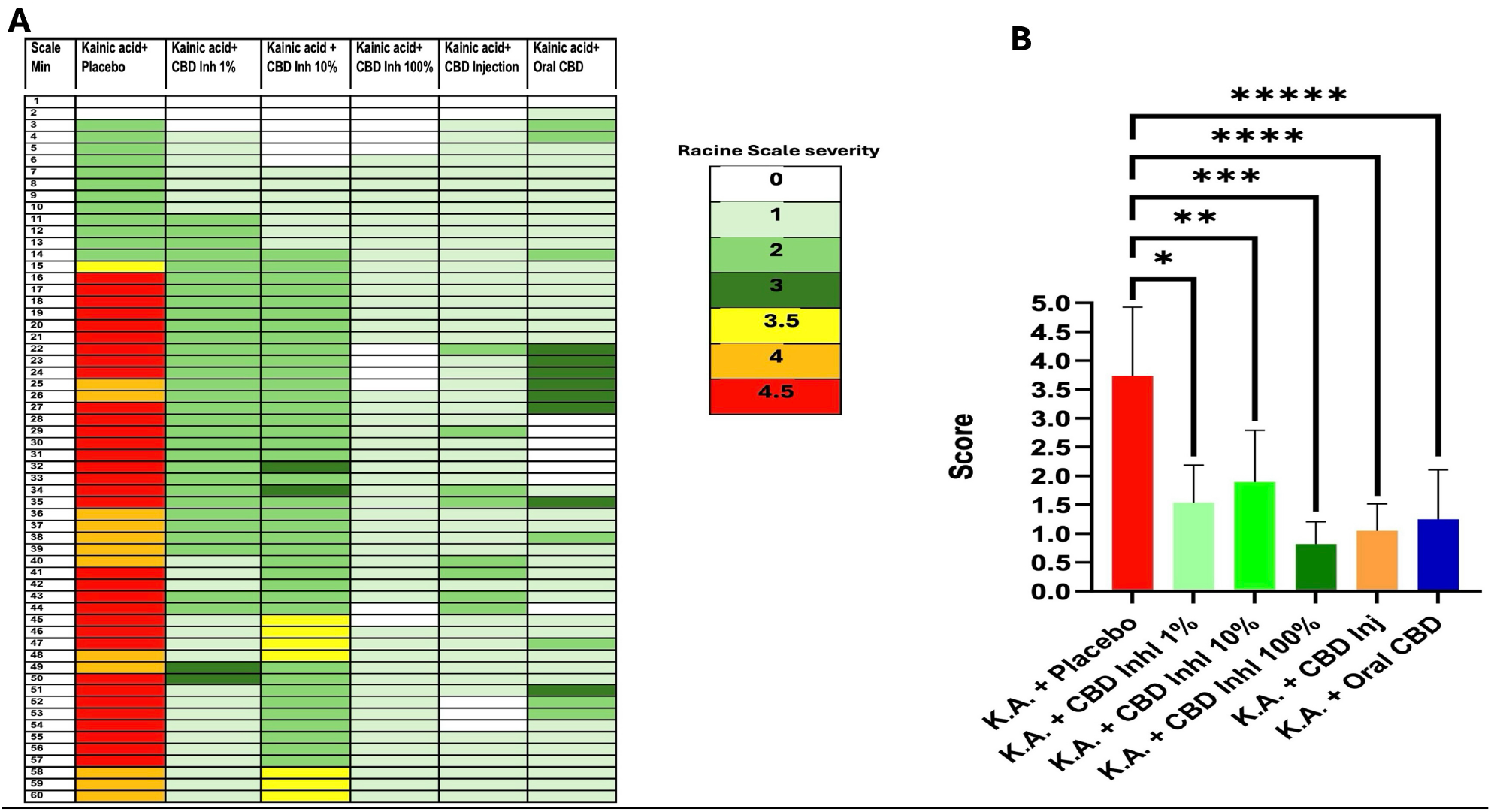
Prophylactic treatment with inhalant CBD reduced severity of kainic acid-induced epileptic seizures. Seizures were induced using 20mg/kg Kainic Acid through intraperitoneal injection to mice. Mice were pre-treated with placebo or one of the variable formulations of CBD including inhaled (3 different concentrations), oral or intraperitoneal injection (i. p.) 30 minutes prior to seizure induction, **a)** Seizure severity was video recorded, scored based on Racine scale ranging from stage 0 to 5, with 0 indicating no abnormality; 1, Mouth and facial movements; (2) Head nodding; (3) Forelimb clonus; (4) Rearing; (5) Rearing and falling, **b)** Quantified histogram of Racine scoring system showing significantly reduced seizures using inhalant CBD compared to placebo group (**p≤0.01, n=5). While other delivery methods of CBD, Oral and i. p. resulted in less seizures, however the difference was not statistically significant (ns, n=5).

### The efficacy of CBD in the treatment of epileptic seizures was route dependent

As shown in Figure 2B, all three forms of CBD used in this study were able to reduce severity of the seizures. However, the inhalation of CBD was the most effective route of delivery in preventing development of symptoms and reducing severity of seizure with a significant difference compared to oral route or i. p. injection (p<0.05). Interestingly, the i. p. route of CBD was more effective in mitigating the seizure severity compared to oral delivery (Fig 2B).

### CBD treatment alleviated the symptoms and reduced severity of epileptic seizures by regulating the expression of neuro-immunologic factors in a route dependent fashion

#### Regulation of alarmin and Immune mediator

Flow cytometry analysis demonstrated that CBD treatment was able to modulate inflammatory responses in both peripheral blood as well as in CNS. CBD decreased the expression level of pro-inflammatory cytokines IL-33, IL-6 in a route dependent manner (Figs 3 & 4 A-C). CBD inhalation showed the most significant measured differences in IL-33 and IL-6 compared to the placebo group (\*\*\*\**p*≤ 0.0001). While oral form of CBD reduced both IL-33 and IL-6 with less significance than inhaled CBD, the injection of CBD lowered the expression of IL-33 and IL-6 with no significant and/or with minimal differences compared to placebo group. Such important and novel findings indicated the significance of route of administration and the outcomes.

**Figure 3.**
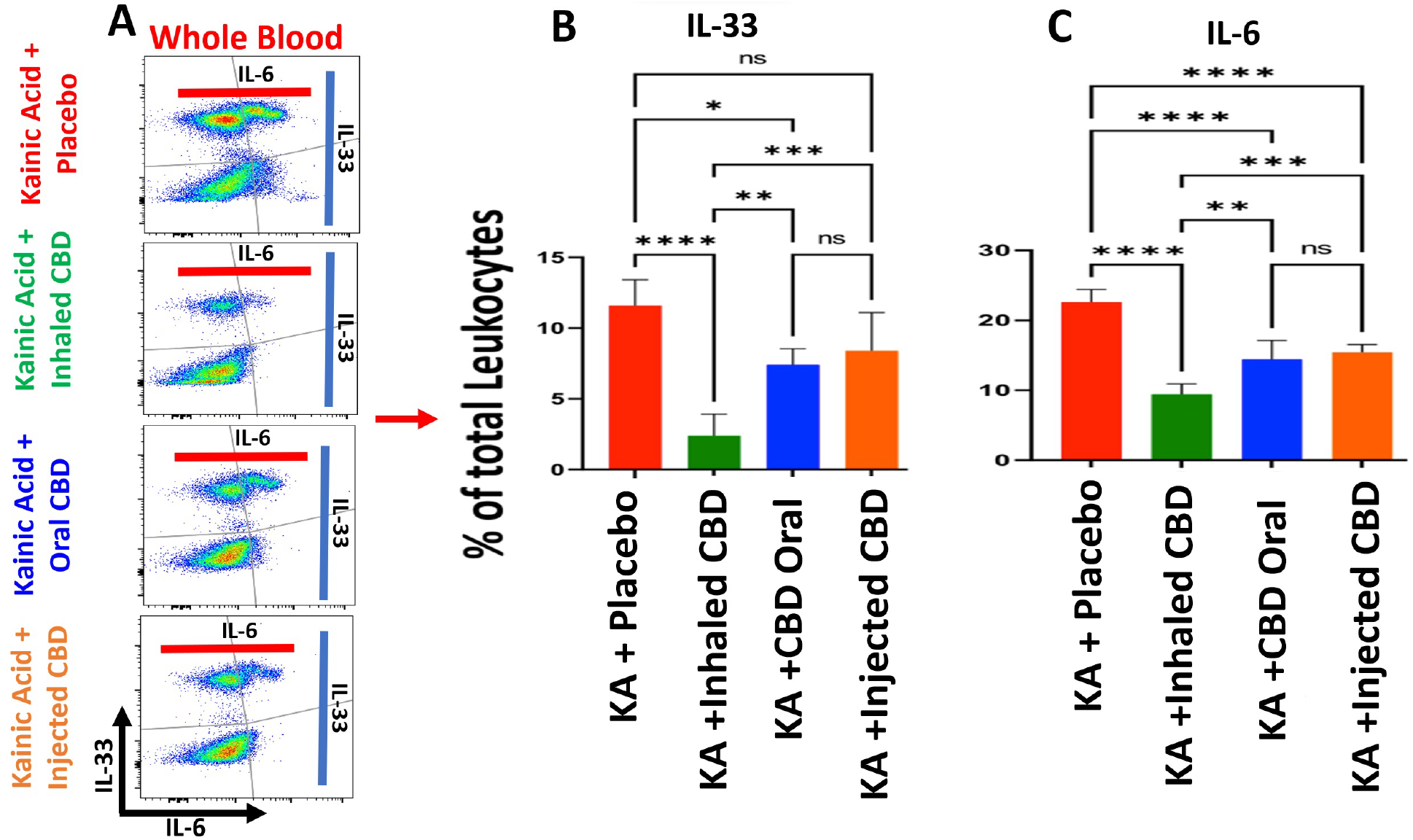
CBD decreased circulating alarmin and pro-inflammatory cytokines in peripheral blood in KA-induced epilepsy. **a)** Flow cytometry of whole blood two-dimension dot plots of IL-33 expressing cells vs IL-6 positive cells in peripheral blood, **b)** Quantified bargraphs of flow cytometry analysis of blood demonstrated a significant reduction in IL-33 level in blood during epilepsy when pre-treated with inhalant CBD, highest reduction (****p≤0.0001, n=5/group) followed by oral form (*p≤0.05, n=5/group). Injected form of CBD delivery, i. p., did not result in any significant reduction of IL-33 in peripheral blood (ns), **c)** Quantified bargraphs of flow cytometry analysis of blood showed a significant reduction in IL-6 in animals during KA-induced epilepsy pre-treated with all forms of CBD compared to placebo (****p≤0.0001, n=5/group). Flow cytometry dot plots are representatives of 5 animals per experimental group.

**Figure 4.**
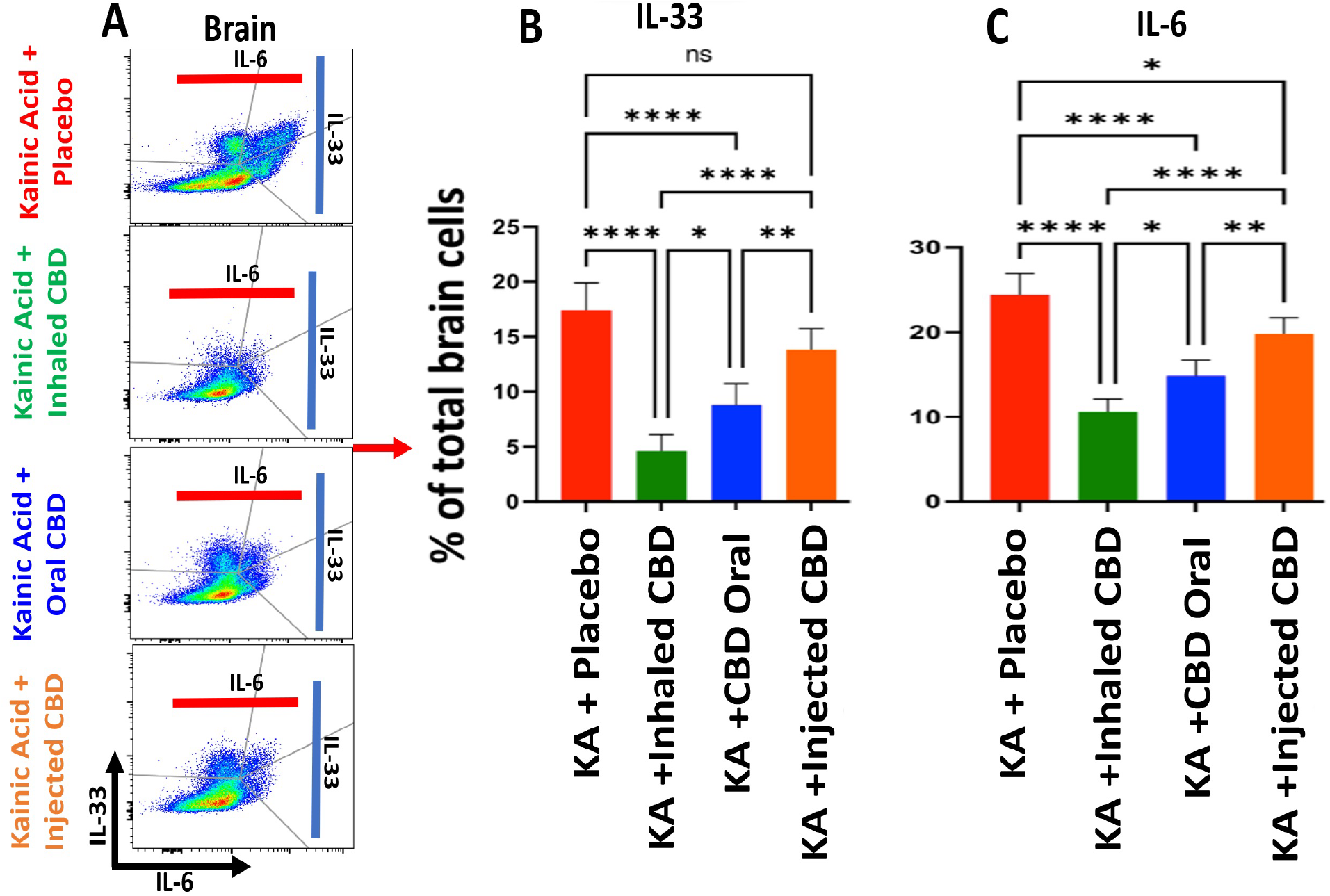
CBD decreased brain alarmin and pro-inflammatory cytokines in KA-induced epilepsy. **a)** Brain flow cytometry two-dimension dot plots of IL-33 expressing cells vs IL-6 positive cells in brain, **b)** Quantified bargraphs of flow cytometry analysis of brain demonstrated a significant reduction in IL-33 level in brain during epilepsy when pre-treated with inhalant CBD or with oral form (****p≤0.0001, n=5/group). Similar to blood, i. p. form of CBD delivery did not result in significant reduction of IL-33 in brain (ns), **c)** Quantified bargraphs of flow cytometry analysis of brain showed a significant reduction in IL-6 in animals during KA-induced epilepsy pre-treated with all forms of CBD. CBD delivery through Inhalation and oral route had identical impact on regulation of IL-6 expression (****p≤0.0001, n=5/group) while the impact of CBD i. p., injection on brain IL-6 was still significant, however with lower level of differences compared to placebo group (*p≤0.05, n=5/group). Flow cytometry dot plots are representatives of 5 animals per experimental group.

#### Alteration of Brain-derived neurotrophic factor (BDNF)

Flow cytometry analysis demonstrated that CBD treatment was able to reduce the expression level of BDNF in both peripheral blood and brain, with the most significant decrease in the group treated with inhaled CBD, followed by i. p. route. Although, the oral administration of CBD showed reduction reduction in the level of BDNF, however, such decrease was not significant and/or minimal compared to placebo group (Figure 5).

**Figure 5.**
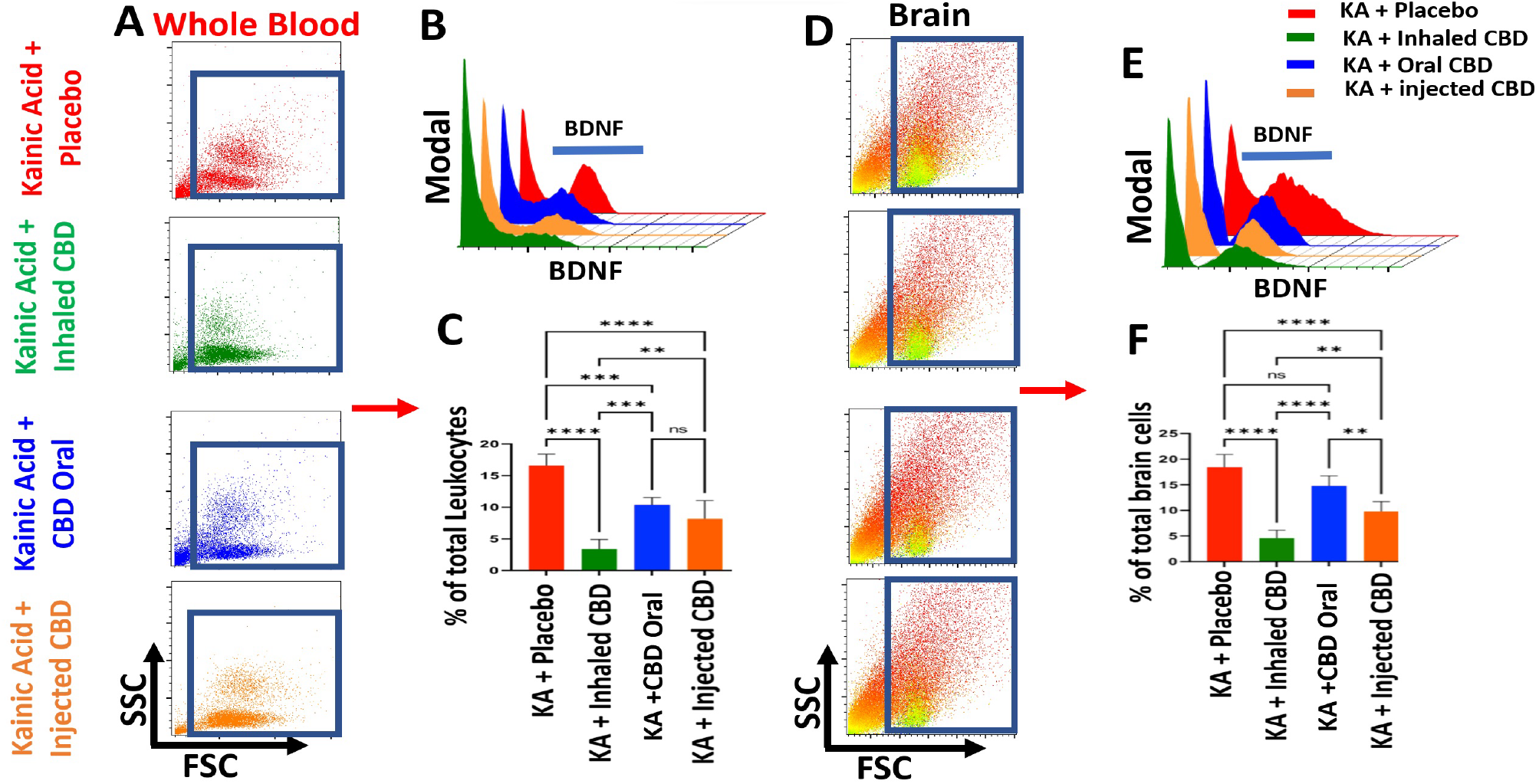
Inhalant CBD reduced BDNF significantly in KA-induced epilepsy. Flow cytometry analysis of peripheral blood demonstrated a significant reduction in BDNF level of mice during epilepsy when pre-treated with CBD. The most reduction in circulating BDNF of blood was when mice pre-treated with either inhalant or i. p. CBD (****p≤0.0001, n-5/group), while oral form resulted in less reduction (***p≤0.001, n=5/group). **a)** Forward scatter vs Side scatter dot plot of whole blood (FSC/SSC), **b)** three-dimension comparative histograms of BDNF in blood of all experimental groups, **c)** quantified bargraphs of blood’s level of BDNF in all experimental groups. For brain tissues, while flow cytometry analysis showed the most significant reduction of brain’s BDNF in mice pre-treated with either inhalant or injected CBD (****p≤0.0001, n-5/group), however, the oral form of CBD did not result in any significant reduction in brain’s BDNF compared to placebo group (ns), **d)** Forward scatter vs Side scatter (FSC/SSC) dot plots of brain tissues, **e)** three-dimension comparative histograms of BDNF in brain tissues of all experimental groups, **f)** quantified bargraphs of brain’s level of BDNF in all experimental groups. Flow cytometry dot plots are representatives of 5 animals per experimental group.

#### Regulation of Immune checkpoints

Flow cytometry analysis demonstrated that CBD was able to up-regulate the expression of Immune checkpoint, Programmed cell death protein-1 (PD-1) in both peripheral blood and in the brain (Fig 6 A-C). Although, CBD in all forms of administration regulated the PD-1, however, it was only inhalant CBD that resulted in a significant difference in PD-1 expression (***p≤0.001) compared to placebo group in both peripheral blood as well as in the brain. Both oral or injected form of CBD administration did not result in a significant difference in the expression level of PD-1 in blood and brain compared to the placebo group.

**Figure 6.**
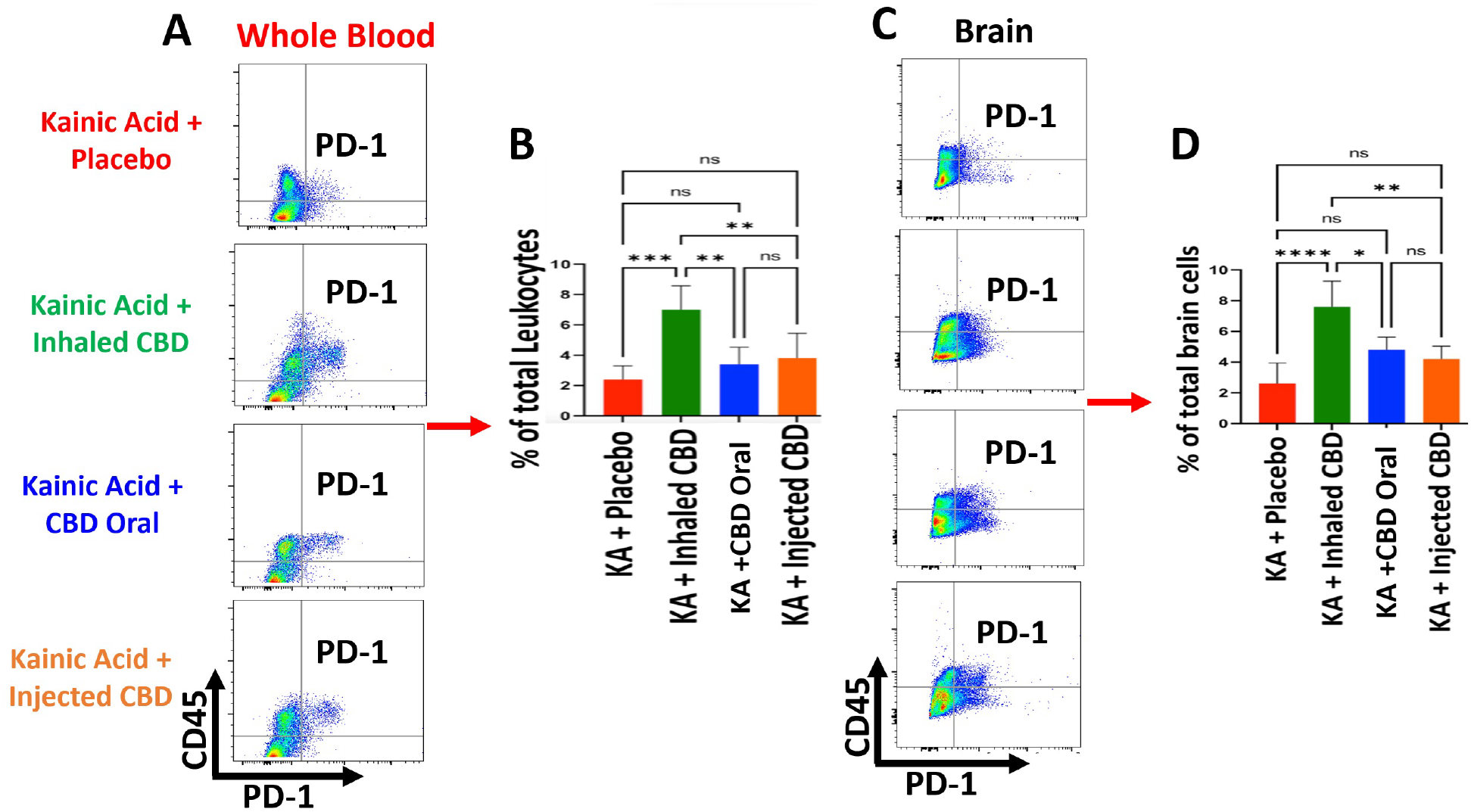
Inhalant CBD increased Immune checkpoint, PD-1, in KA-induced epilepsy. Flow cytometry analysis of peripheral blood demonstrated a significant elevation in circulating PD-1 level of mice in blood during epilepsy when pre-treated with inhalant CBD (***p≤0.001, n-5/group). Oral form or injected form of CBD (i.p) did not result in any significant increase of PD-1 (ns), **a)** Whole blood flow cytometry two-dimension dot plots of leukocytes (CD45+ cells) vs PD-1 expressing cells in the whole blood, **b)** quantified bargraphs of blood’s level of PD-1 in all experimental groups. For brain tissues, while flow cytometry analysis showed a significant increase in brain’s PD-1 in mice pre-treated with inhalant CBD (****p≤0.0001, n=5/group), however, the i. p., or oral form of CBD did not result in any significant increases in brain’s PD-1 compared to placebo group (ns), **c)** Brain tissue’s two-dimension flow cytometry dot plots of microglia/macrophages (CD45+ cells) vs PD-1 expressing cells in brain tissues, d) quantified bargraphs of brain’s level of PD-1 in all experimental groups. Flow cytometry dot plots are representatives of 5 animals per experimental group.

## Discussion

Our findings demonstrate that CBD pre-treatment reduced the severity of the epileptic seizures in an experimental model. Most importantly, our data showed, for the first time, that CBD efficacy could be route- and type-dependent, with inhaled CBD broad spectrum resulting in the most substantial reduction in the severity of the seizures and limiting the symptoms compared to other routes of administration (ROA) and type of CBD. Given the crucial role of ROA, such novel findings could be a major determinant in the therapeutic processes to ensure that medications are delivered to the targeted sites accurately and effectively (Strang et al; 1998; Ruiz and Montoto, 2018). ROA can influence drug bioavailability, affecting local and systemic absorption, contributing to the speed of action, latency, and intensity of medications (28-29). Importantly, a desirable ROA should be patient friendly with minimal side effects in a non-invasive fashion (28-29). Although, several reports have described advantages and disadvantages of common ROAs for CBD (oral, sublingual, topical, inhalant, intranasal, rectal, and parenteral) individually (30-31), to date, no conclusive studies have investigated features and outcomes of ROAs in a comparative fashion specifically regarding epileptic seizures with CBD treatment. Our findings in this study suggest not only inhaled CBD as a novel treatment for epileptic seizures, but also provide evidence for superiority of inhaled CBD compared to other ROAs including oral and intraperitoneal injection of CBD with significant differences (p<0.05). The first pass effect is one of the factors, which reduces the efficacy of the drug especially when it is administered through oral route. As a result of the first pass effect phenomenon, the concentration and efficacy of the drug would be significantly reduced before reaching the main target. The inhalation route, however, not only accelerates delivery time and accuracy, but it also improves the efficacy by minimizing the first pass effect, enhancing efficacy in a shorter period. Furthermore, inhalers use a lower dose compared to the oral route in safer and more cost-effective fashion. Other advantages of inhaled therapies compared to other routes of administrations include higher efficacy, less invasive, faster absorption with higher bioavailability and fewer systemic side effects.

Importantly, our data showed, for the first time, that CBD treatment regulated immunologic responses to seizures by lowering proinflammatory cytokines (IL-33 and IL-6) production and enhancing the level of Immunecheckpoint (IC) protein of PD1 within the CNS as well as systemically in the peripheral blood. Increasing evidence points to inflammation and oxidative stress as two major contributing factors in the formation, permanence, and recurrence of seizures (32-33). Cytokines and their receptors are major mediators and modulators of inflammatory responses within the CNS and peripheral tissues. In fact, the impact of cytokines’ functions within the CNS goes beyond immunologic modulation, affecting neurons acting as neuromodulators as described by several studies (34). Interleukin 33 (IL-33) is a pleiotropic cytokine and a member of the IL-1 family which is released as an alarm signal (alarmin), as part of pattern-recognition system: DAMP (Damage-Associated Molecular Pattern), from apoptosis and necrosis when cells are under stress conditions (35). Therefore, it is logical to target IL-33 as a neuro-immunotherapeutic modality in the treatment of epileptic seizures (36). Nevertheless, several studies have suggested a dichotomic function of IL-33 during neuro-inflammatory responses (37-38). Such dual action means that the impact of IL-33 could be affected by several factors including, not limited to, epigenetic and micro-environmental variables. While such binary action of IL-33 magnifies its central role in the inflammatory responses within the CNS, it also accentuates the significance of CBD as a key regulator, not a suppressor, of IL-33 in optimal and timely modulation of IL-33. A recent report from our group showed a role for CBD-induced regulation of IL-33 in a murine model of Alzheimer’s disease (8). Understanding the mechanisms by which CBD influences the IL-33 shift from a damaging pro-inflammatory alarmin to a protective cytokine with therapeutic values requires further research and elucidation. IL-6 is also a pro-inflammatory cytokine with convulsive nature contributing to the progression and severity of epileptic seizures (F39-40). Several studies have shown increased level of IL-6 during inflammation within the CNS domain significantly higher than homeostasis and normal condition (Jia et al; 2020). Our findings here and in previous studies, have shown that CBD can regulate IL-6 effectively in different pathologic conditions (41). Such potential could be further investigated as an innovative and effective therapeutic modality in the treatment of a wide spectrum of inflammatory diseases such as epilepsy and its complications.

One of the major immunologic molecules in regulation of cytokines and inflammatory mediators is Immune checkpoints (ICs). Immune checkpoints are gatekeepers of Immune responses, restoring and maintaining the immune balance through modulation of innate and adaptive effectors of immune system (42). Programmed cell death protein 1 (PD-1), is considered as a major IC molecule expressed mainly on T-cells as well as several other types of Immune cells including B cells, NK cells, macrophages, and dendritic cells (43). The role of PD-1 in malignancies and auto-immune diseases is well established (44). Increasing evidence supports the notion that PD-1 can influence the maintenance and re-establishment of homeostasis in CNS during health and in diseases (45). However, the role of PD-1 in epilepsy has not been fully understood (46). Few studies have reported the effects of PD-1 in epilepsy or proposed the use of PD-1 as a biomarker in the early diagnosis as well as a therapeutic target in the treatment of epilepsy and seizure prevention (46). Our findings support the notion of the role of PD-1 in epilepsy and seizure severity. In addition, based on our novel findings, we are further proposing that CBD has the potential to control and ameliorate seizure severity through regulation of PD-1. Inhalant CBD could be a non-invasive Immuno-therapeutic modality to contain excessive inflammation by up-regulation of ICs, restoring the immune balance and homeostasis. Such potential could have implications in several other neurologic and inflammatory disorders.

Furthermore, our results demonstrate that induction of epilepsy enhanced the level of brain-derived neurotrophic factor (BDNF) systemically in blood and locally within brain tissues. This was an expected observation since several studies have already demonstrated the link between BDNF and epilepsy (47). However, the novel findings in this study showed the potential of CBD in the regulation of BDNF in our model of seizure induction. There could be several hypothetical theories to explain the link between CBD and BDNF. It is already reported that it could be a potential connection between cannabinoids and a receptor for BDNF, Tropomyosin receptor kinase B (TrkB) (48). Therefore, it is possible that CBD may affect the BDNF/TrKB signaling and influence the severity of epileptic seizures, which necessitates further investigation.

Additionally, one of the most important findings of this study presented is the central role of CBD formulation. While current CBD-based treatment for epileptic seizure uses CBD isolate, our findings suggest that a formulation with broad spectrum CBD may have more beneficial impact on the seizure resolution. This is consistent with the notion of “entourage effect” and higher efficacy in the disease resolution based on the superior function of cannabinoids in a synergistic dynamic of formulation (49). This means that the integrality of cannabis product, including active and inactive components is more important and effective than only the active products by themselves (49). Our discovery necessitates the re-examination and assessment of current CBD formulations in the treatment of seizures and epilepsy.

In conclusion, while CBD has already been shown to have beneficial effects on epileptic seizures, the efficacy of current CBD-based treatments needs optimization by elucidation of mechanism of action, ROA, and formulation. Our findings suggest that route of administration and CBD formulation play crucial roles in the efficacy of treatment. CBD delivery through inhalation appears to be more effective in reducing the seizure severity in a non-invasive fashion compared to other existing CBD-based medications for epileptic seizures. In addition to being more effective, the cost of inhalant CBD is also much more affordable than either injected or oral formulations. The use of isolate versus other formulations of CBD such as broad-spectrum CBD seems to affect treatment efficacy, requires, and warrants further and extensive research.

## Limitations

Our novel findings in this study were associated with some limitations. Our study was performed to assess the preventive effects of inhaled CBD in a prophylactic fashion. Further studies are required to demonstrate the efficacy of inhaled CBD in the treatment of seizures in a post-induction mode. Further, while the cellular and molecular alterations reported during these studies are highly novel, however, to determine the causality of CBD effects on such changes remain to be further studied.

## Abbreviations

ASM: Anti-seizure medications
BDNF: Brain-derived neurotrophic factor
CBD: Cannabidiol
CNS: Central Nervous System
DS: Dravet syndrome
FDA: Food and Drug Administration
IC: Immunecheckpoint
IL-33: Interleukin 33
IL-6: Interleukin 6
i.p.: intraperitoneal injection
KA: Kainic Acid
LGS: Lennox-Gastaut syndrome
PD1: Programmed cell death protein 1
ROA: Route of Administration
THC: delta-9-tetrahydrocannabinol
TrkB: Tropomyosin receptor kinase B.

## Data availability statement

All data supporting the findings of this study are available within the paper and also are available upon request from the corresponding author.

## Acknowledgement

Authors are thankful to ThriftMaster Holding Group for providing the inhalant CBD for this study. Authors also thank Medicinal Cannabis of Georgia for providing help in optimizing the CBD dosage.

## Ethical publication statement

We (authors) confirm that we have read the Journal’s position on issues involved in ethical publication and affirm that this report is consistent with those guidelines.

## Author contributions

All authors contributed to the study, commented on the manuscript, and approved the final version

